# Confirmation of Highly Pathogenic Avian Influenza (HPAI) H5N1 Associated with an Unexpected Mortality Event in South Polar Skuas (Stercorarius maccormicki) during 2023-2024 Surveillance Activities in Antarctica

**DOI:** 10.1101/2024.04.10.588951

**Authors:** Benjamín Bennett Lazo, Bárbara Berazay, Gabriela Muñoz, Naomi Ariyama, Nikita Enciso, Christina Braun, Lucas Krüger, Miloš Barták, Marcelo González-Aravena, Victor Neira

## Abstract

From December 2023 to March 2024, a surveillance program to detect HPAI H5N1 was performed on Antarctica Territory, at Fildes Peninsula (King George Island, Maritime Antarctic), and James Ross Island. At Fildes Peninsula, samples from marine birds and mammals from four sampling locations were collected, based on their accessibility and the presence of animal colonies: Ardley Island, which presents a high concentration of Gentoo penguins (*Pygoscelis papua*); Ardley Cove were small groups of likely non-reproductive Chinstrap penguins (*Pygoscelis antarcticus*); Southern elephant (*Mirounga leonina*) and Weddell (*Leptonycotes wedellii*) seals haul-out sites; and, a nesting site of Southern giant petrels (*Macronectes giganteus*). On February 28^th^ the surveillance group received a request for sample collection, due to the observation of deceased seabirds at James Ross Island in the area neighboring the Lachman lakes (63.7989 S, 57.8105 W). Six samples from five dead South polar skuas (*Stercorarius maccormicki*) were collected on March 3^rd^, 2024. All samples were collected adhering to the Antarctic Treaty guidelines. After collecting a total of 943 samples from Fildes Peninsula all results tested negative, and no individuals had clinical signs or behaviors compatible with HPAI. However, all samples from South polar skuas from James Ross Island were confirmed positive for HPAI H5N1 clade 2.3.4.4 by specific real-time RT-PCR reactions. These results confirmed the first mortality event registered in Antarctica (south of 60°S) caused by HPAI H5N1, in this case on South polar skuas. Further studies are needed to genetically characterize the virus and to better understand the role of skuas in the dynamics of viral dissemination in Antarctica.

## Introduction

Highly pathogenic avian influenza (HPAI) subtype H5NX, clade 2.3.4.4b has been responsible for several outbreaks worldwide causing multiple mass mortality events among avian wildlife and marine mammal populations (Charostad et al., 2023). The wide and fast dispersion of this virus is attributed to the contamination of migratory species (Caliendo et al., 2022a; Prosser et al., 2022; Gass Jr et al., 2023) and those with long-range dispersing capabilities (Boulinier, 2023; Lane et al., n.d.). On the other hand, the H5 highly pathogenic strains have undergone extensive evolution. The current strain, HPAI H5N1 clade 2.3.4.4b arose in 2020. The most recent major outbreak occurred in the South American sub-continent, with the virus first confirmed in October 2022 (Ariyama et al., 2023). Until then, the virus had not reached Oceania (Australia and New Zealand) nor the Antarctic continent.

Considering the proximity of South America to the Antarctic continent, the migration, and the movements of many seabird species between South America and the Antarctic Peninsula throughout the year, several concerns have arisen about the possible arrival of the virus along with animals moving from positive HPAI locations. Previous evidence strongly suggests the dissemination of the Influenza virus to the Antarctic continent, where the detected viruses are genetically correlated to those detected outside Antarctica (Barriga et al., 2016; de Seixas et al., 2022). In addition, there is also evidence that Arctic terns *Sterna paradisaea* (Egevang et al., 2010) and South polar skuas *Stercorarius maccormicki* (Kopp et al., 2011) can do trans-equatorial migrations from and to the Antarctic continent, both of which have been proven to be capable of viral shedding.

However, since the arrival of the virus to the southern hemisphere a more relevant role may played by susceptible animals with shorter migration patterns and/or wide foraging areas like giant petrels (*Macronectes* spp.), Brown skuas (*Stercorarius antarcticus lonnbergi*) (Krietsch et al., 2017), and even Southern elephant seals *Mirounga leonina* (Hindell et al., 2016), that may come in contact with infected individuals, coming back to Antarctic regions before developing clinical signs.

Although some positive cases of HPAI H5NX 2.3.4.4b. have been confirmed in South Georgia island, considered a Sub-Antarctic territory (BAS, 2024), there are no publications confirming cases in southerly latitudes. This fact may suggest that Antarctic species may still be largely susceptible to the infection, being vulnerable to high mortality events in case of the arrival of the virus in Antarctic animal colonies (Dewar et al., 2023). In this context, we performed active surveillance to identify HPAI in seabirds and marine mammals in several locations of King George Island, South Shetland Islands, Maritime Antarctic Peninsula. This area may play a substantial role as an entry point to the Antarctic continent due to its proximity to locations previously reported as positive and, more importantly, there is an increased proximity between wildlife and humans (Pertierra et al., 2017; McCarthy et al., 2022), which may facilitate viral dissemination. In addition, we were in contact with researchers in other localities of the Antarctic Peninsula to communicate with us in case they observed unexpected mortality events during their activities so we could take samples, or also receive samples from other locations. Thus, we received a request from investigators of the Czech Antarctic research team, due to the observation of deceased seabirds 4 km East of the Mendel Base at James Ross Island (skua nesting site close to the Lachman lakes (63.7989 S, 57.8105 W). This document resumes the results obtained during December 2023 and March 2024 in an active surveillance program confirming the presence of HPAI H5N1.

## Materials and Methods

### Locations at Fildes Peninsula, King George Island

The surveillance team was based at Professor Julio Escudero Base, King George Island, South Shetland Islands, Maritime Antarctic. Between 16 December 2023 and March 23^rd^, 2024, frequent surveillance expeditions were carried out in the proximity of the base (see Fig 1) covering the hotspots of wildlife including penguins, flying seabirds, and marine mammals. In those expeditions, clinical observation and sample collection were performed. The locations present animal aggregations and colony sites (Braun et al., 2012). Thus, 10 locations were selected to visit at least once (Fig 1c): Ardley Cove in front of Professor Julio Escudero and Bellingshausen research stations, where Gentoo (*Pygoscelis papua*) and Chinstrap (*Pygoscelis antarcticus*) penguins are often seen resting; the Ardley Island, Antarctic Specially Protected Area (ASPA 150), where there are small breeding groups of Adélie penguins (*Pygoscelis adeliae*) and a large Gentoo penguin breeding colony; Hydrographers Cove, where non-breeding chinstrap penguins and skuas are seen with hauling-out Weddell seals (*Leptonychotes weddellii*); Diomedea Island with a breeding colony of Southern giant petrels (*Macronectes giganteus*) with 62 breeding pairs; Flat-top Peninsula, where Weddell and Elephant seals haul-out are found; Biologists Cove, also an important site for Southern elephant, Weddell and Antarctic fur (*Arctocephalus gazella*) seals; Basalt Creek beach line, also an important site for Elephant, Weddell and Fur seals where chinstrap penguins and Antarctic shags breed; Gradzinski Cove, where Elephant and Weddell seals haul-out, a small breeding colony of Antarctic Fur seals can be found nearby an old shelter, and on the upper terrains several nests of Southern giant petrels (177) can be found; and Green Point, where numerous Kelp gulls (*Larus dominicanus*) nest. The complete list of locations is detailed in Table 1.

**Figure 1.**
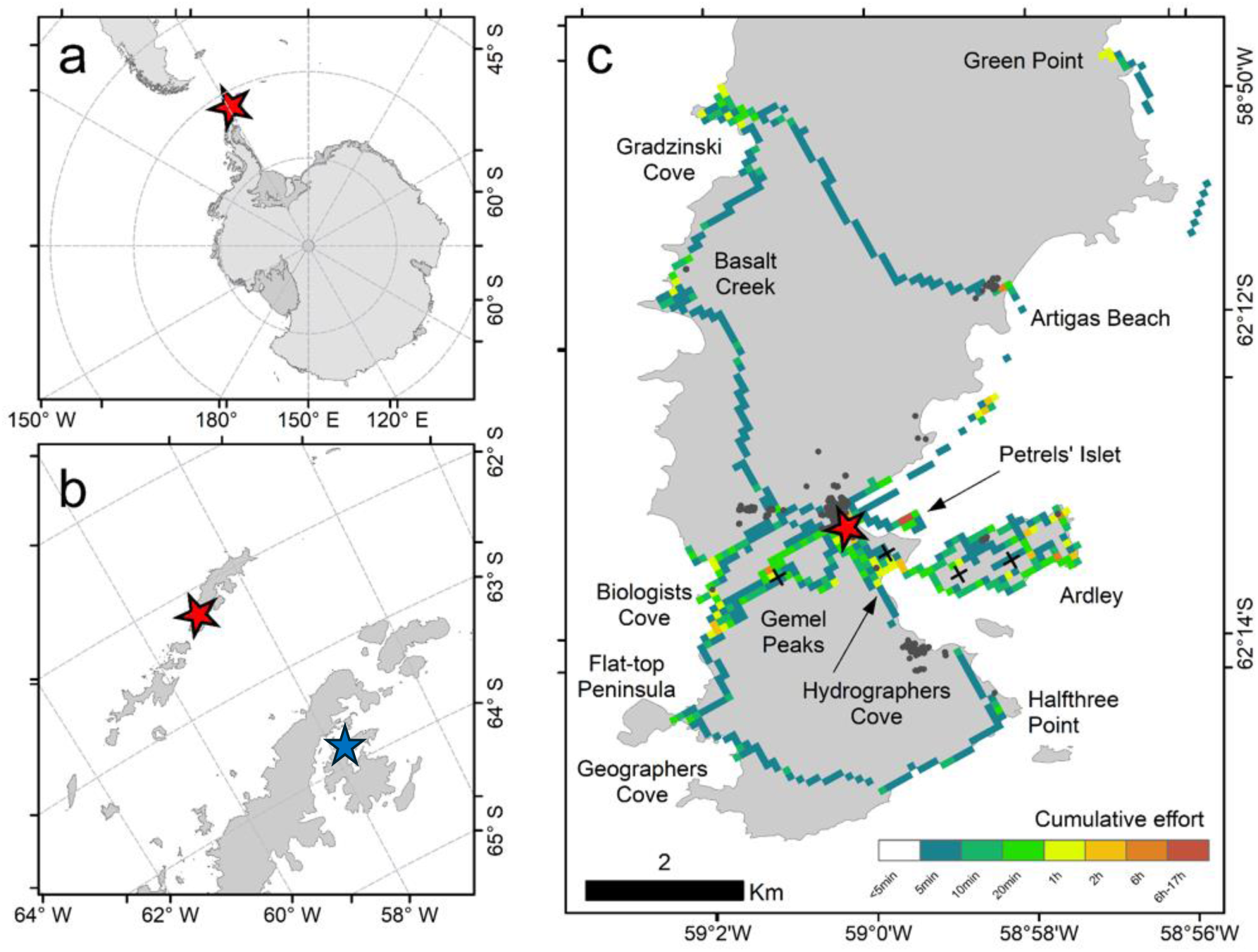
Location of the Escudero Research Station (red star) and James Ross Island (blue star) on the scale of the Antarctic Continent (a), on the scale of the South Shetland Islands (b) and Fildes Peninsula in detail (c) with the cumulative effort calculated from tracking researcher movements during sampling and surveillance for symptomatic animals with clinical signs compatible with high pathogenic avian influenza. Names of locations in ‘c’ from the Composite Gazetteer of Antarctica, Secretariat SCAR https://data.aad.gov.au/aadc/gaz/scar/download.cfm. Seal samples were taken in Hydrographers Cove, Flat-top Peninsula, Biologists Cove, Basalt Creek, and Gradzinski Cove; penguin samples were taken from Ardley Cove, Ardley Island, and Hydrographers Cove; giant petrel samples were taken from Diomedea Island (Petrels’ Islet); breeding shag samples were taken in Ardley Island; Black ‘x’ are location of Skuas’ samples in Ardley Island, Hydrographers Cove, and Gemel Peaks. See Table 1 for details on samples taken. Further fauna observations (without sampling) were done through every covered area, emphasizing 78 Southern Giant Petrel active nests in the upper terrains of Gradzinski Cove and 52 active nests in Diomedea Island; two skua nests between Halfthree Point and Geographers Cove and two nests in the upper terrains of Gradzinski Cove; Elephant, Weddell, and Fur Seals in Geographers Cove, Flat-top Peninsula, Green Point, and a small group of breeding Fur Seals in Gradzinski Cove; one Leopard Seal and Breeding Kelp Gulls in Green Point; Chinstrap Penguins in Ardley Cove, Hydrographers Cove, and Basalt Creek.

**Table 1.** Sampling details during high pathogenic avian influenza (HPAI) surveillance Sampled locations, species, types of samples (environmental feces swabs and direct cloacal swabs), and quantities. Positive samples were collected only from James Ross Island.

| Locations | Species | Environmental | Direct | Total |
| --- | --- | --- | --- | --- |
| **Ardley Island** | Gentoo penguin | 725 (145 pools of 5 fecal samples) | 2 (dead brain sample) | 727 |
|  | Adelie penguin | 40 (8 pools 5 fecal samples) | - | 40 |
|  | Antarctic Shag | - | 4 (cloacal swab) | 4 |
|  | Chinstrap Penguin | - | 3 (cloacal swab) | 3 |
|  | Brown Skua | 2 | 1 | 3 |
| **Hydrographers Bay, King George Island** | Gentoo penguin | - | 2 | 2 |
|  | Weddell seal | 8 | - | 8 |
|  | Chinstrap penguin | - | 9 (cloacal swab) | 9 |
|  | Brown Skua | - | 1 (brain sample) | 1 |
|  | Emperor penguin | - | 1 (cloacal swab) | 1 |
| **Gemel peaks** | Brown Skua | - | 2 (cloacal swab) | 2 |
| **Diomedea Island (Petrel's Islet)** | Southern Giant Petrel | - | 39 (cloacal swab) | 39 |
|  | Unidentified | 1 | - | 1 |
| **Elephant's beach** | Chinstrap Penguin | - | 23 (brain and lung samples) | 23 |
|  | Elephant seal | 29 | - | 29 |
|  | Weddell seal | 4 | - | 4 |
|  | Fur seal | 2 | - | 2 |
|  | Gentoo penguin | 5 (1 pool 5 of fecal samples) | - | 5 |
|  | Unidentified | 1 | - | 1 |
| **Basalt Creek and Gradzinski Cove** | Elephant seal | 11 | - | 11 |
|  | Weddell seal | 4 | - | 4 |
|  | Fur seal | 3 | - | 3 |
| **Ardley Cove** | Chinstrap penguin | 5 (1 pool 5 of fecal samples) | - | 5 |
|  | Gentoo penguin | 5 (1 pool 5 of fecal samples) | 1 (cloacal swab) | 6 |
| **Collins Base** | Antarctic gull | 10 | - | 10 |
| **James Ross Island** | Polar Skua | - | 6 (tissue pool samples) | 6 |
| **Total** |  | 857 | 94 | 949 |

Sampling effort was recorded by tracking researchers’ movements during fieldwork using a Garmin InReach Explorer and a Garmin Enduro Watch in tracking mode to record geographic positions every 60 and 30 seconds respectively. The tracking data was processed using ‘trip’ R-package (Sumner et al., 2009) to quantify time spent at different sites, and ArcMap was used for mapping the distribution of effort. In addition, the clinical observations (search of respiratory, neurologic, or digestive syndromes) were conducted by veterinarians. Animal surveillance was performed in all groups of the visited locations, monitoring sudden animal deaths or individuals with clinical signs compatible with HPAI H5N1. The study was approved by the Institutional Ethics Committee of the University of Chile number 22603-VET-UCH.

### Sampling

Sampling activities were conducted under permission numbers 669-2023, 670-2023, and 201-2024 by INACH. During each visit, researchers looked for evidence of disease and collected environmental and direct samples. In general, the sampling was focused on collecting fresh fecal/dropping samples to minimize animal handling and maximize the number of samples. Direct samples were obtained from live Southern giant petrels, skuas, and opportunistic such as in fresh dead animals. Samples were collected using rayon swabs, which were deposited in Viral Transport Media (Inactivated Type) (ALLTEST, catalog number ITM-001). In the case of environmental samples, they were collected and pooled in groups of 5 samples into one tube of viral transport media. In the case of individual cloacal swabs, the individuals were captured and restrained following pre-approved protocol numbers. 069/CEC/2018 and 3/CBSCUA/2022. All samples were collected following Antarctic Treaty guidelines by a surveillance group composed of a biologist and four veterinarians, who were also responsible for the observation of clinical signs compatible with the disease.

### Unexpected mortality case at James Ross Island

On February 28^th^, 2024, we received a report of three dead skuas (unidentified species) on James Ross Island, 63.7989 S, 57.8105 W noted by the field crew of the Czech Antarctic Research Program near the Base Johann Gregor Mendel (Figure 1b). The researchers, who were not related to animal research but with 20 years of experience in the location, indicated that three dead individuals in one spot were never recorded from this place. The site is a typical nesting area at the NE tip of James Ross Island (Weidinger et Pavel 2013) and the spot where skuas wash in the freshwater of two large lakes and several small-area freshwater ponds. Our surveillance team traveled from King George’s Island to James Ross Island in the Chilean vessel Janequeo (ATF-65) arriving on March 3^rd^, 2024. At the place, five deceased animals were found and identified as South polar skuas (Figure 2), samples were collected from all individual animals, but in pools from different tissues including the brain (Table 2). Sampling was conducted following the protocol previously mentioned.

**Figure 2.**
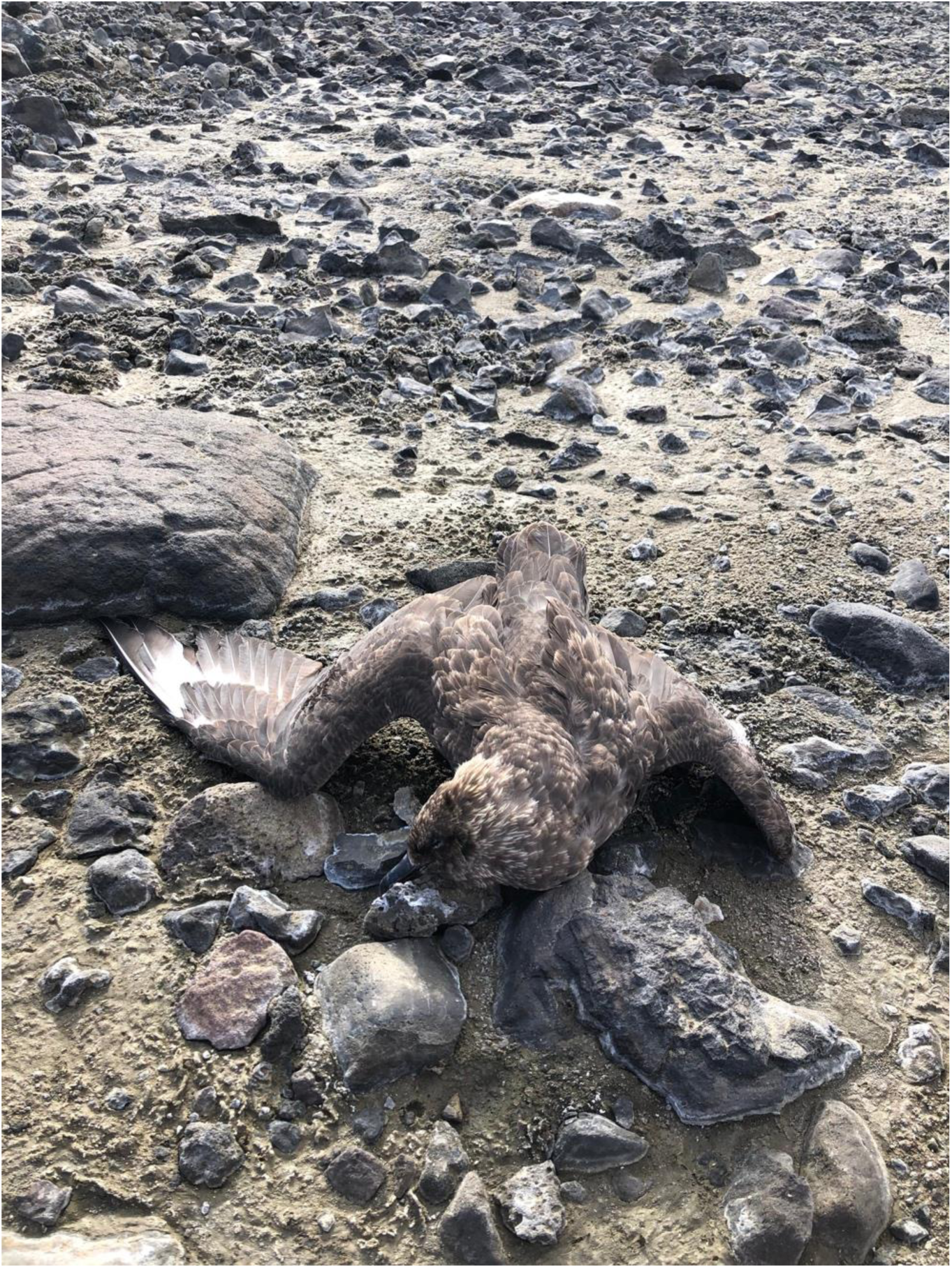
Deceased South Polar Skua at James Ross Island.

**Table 2.** Real-time RT-PCR results from suspected South Polar Skua at James Ross Island. All samples were confirmed positive for HPAI H5N1 2.3.4.4.

| ID | Sample | Species | Gene M | H5 2.3.4.4 | HPAI* | N1 | Interpretation |
| --- | --- | --- | --- | --- | --- | --- | --- |
| 1 | Brain-Lung | South Polar Skua | 16.4 | 15.4 | 13 | 16.6 | HPAI H5N1 2.3.4.4 |
| 2 | Brain-Lung | South Polar Skua | 18.6 | 18 | 16.1 | 18.8 | HPAI H5N1 2.3.4.4 |
| 3 | Brain-Lung | South Polar Skua | 19.4 | 18.3 | 17.5 | 20.7 | HPAI H5N1 2.3.4.4 |
| 4 | Brain | South Polar Skua | 18.8 | 17.6 | 15.8 | 18.7 | HPAI H5N1 2.3.4.4 |
| 5 | Brain-Trachea | South Polar Skua | 16.7 | 15.5 | 13.9 | 16.5 | HPAI H5N1 2.3.4.4 |

\* HPAI pathotyping

### Diagnostic testing

The samples were processed within 24 hours after collection at Professor Julio Escudero base’s molecular biology laboratory. The RNA from the samples was extracted using the acid guanidinium thiocyanate-phenol-chloroform method with Chomczynski phenol solution (Winkler Ltda.) following the manufacturer’s recommendations. Real-time Retro-Transcriptase Polymerase Chain Reactions (RT PCR) were performed, amplifying the conserved region of the M gene of general Influenza virus (WHO, 2011). Samples over a CT value of 35 were considered inconclusive and over a CT value of 40 were considered negative for Influenza virus and, therefore negative for HPAI. The positive samples were subtyped and pathotyped to confirm the HPAI H5N1 clade 2.3.4.4 using the NVSL-specific protocols for H5 clade 2.3.4.4, H5 pathotyping, and N1 subtype (1732.02, 1767.01, and 1768.01) (Ulloa et al., 2023; Ariyama et al., 2023).

## Results

### Effort

On the Fildes Peninsula, our observation and sampling efforts reached a total of 141 hours and 30 minutes, covering at least 209.5 km of transections (Figure 1). Higher observation effort (Figure 1) was concentrated in Diomedea Island (25.4 hours), Ardley Island (22.7 hours), Biologists Cove (10.8 hours), Gemel Peaks (9.2 hours), and Gradzinski Cove (7.4 hours).

### Lack of HPAI in King George Island

Overall, we observed breeding and non-breeding Gentoo, Chinstrap, and Adélie penguins, Southern giant petrels, Antarctic shags (*Leucocarbo bransfieldensis*), Antarctic tern (*Sterna vittata*), Brown and South polar skuas, Kelp gulls, Antarctic fur seals, non-breeding Southern elephant, Weddell, and Leopard (*Hydrurga leptonyx*) seals (Fig 1c). Not a single observed animal had clinical signs attributable to HPAI. No cases of unexpected mortality events were observed either. The researchers observed a decline in the skua populations at Fildes Peninsula (including Ardley Island), a recognized nesting area, compared to prior years. This observation was more evident for South polar skua than Brown skua. In addition, one fresh skua carcass (non-identifiable) was found near the Ardley Island isthmus, and a few carcasses of Gentoo penguin chicks were found on Ardley Island, which is considered usual mortality. In late February, we observed 15 carcasses of Chinstrap penguins at Elephant’s Beach negative for HPAIV. 943 samples were collected during the surveillance period, of which 857 were environmental fecal samples and 88 direct samples, 58 were cloacal swabs, and 26 were samples from freshly found carcasses. All samples resulted negative for Influenza virus A, reporting no CT value to real-time RT-PCR.

### HPAI H5N1 in South polar skuas

The results of real-time RT-PCR testing on South polar skuas from James Ross Island are presented in Table 2. All the deceased South polar skuas tested positive for HPAIV H5N1 2.3.4.4, with Ct values ranging from 13 to 20.7. There were no signs of predation or scavenging on the animals. Additionally, we did not have permission to collect tissue samples for histopathological examination, and due to our lack of expertise in pathology, we were unable to identify any macroscopic lesions. Notably, two healthy skuas were observed in proximity to the deceased birds.

## Discussion

During the spring of 2022 and summer of 2023 in the southern hemisphere, HPAI was confirmed in most of the countries in South America, then the virus was approaching the Antarctic continent (Charostad et al., 2023). Before the surveillance expedition, the virus was confirmed in sub-Antarctic regions of South Georgia and the Falkland Islands (Bennison et al., 2023; BAS, 2024). Then, this study was designed to monitor the presence of HPAI in marine mammals and seabirds in Antarctica, with a particular focus on the Fildes Peninsula, at King George Island an important biodiverse area where research activities and tourist operations are concentrated (Convey and Peck, 2019). Additionally, we were available to visit other locations if mortality cases were informed.

The HPAI H5N1 virus was confirmed in the mortality event observed on February 28^th^, involving South polar skuas at James Ross Island. The specific TaqMan PCR assays allowed us to precisely confirm the presence of the H5 subtype 2.3.4.4 clade, the highly pathogenic pathotype, and the N1 subtype. While further sequencing and genetic analysis are necessary, it is crucial to report these initial results promptly. The reliability of the multiple PCR reactions gives us high confidence in these findings.

The case was considered a massive and unexpected mortality event, especially since dead Skuas are rarely observed. The cause of death in these animals was most likely the HPAI H5N1 virus because of the low Ct values observed in the samples. This event aligns with other recent information that confirms the virus in skuas in Antarctic territory. On February 25^th^, 2024, a couple of days, before skuas mortality was reported in James Ross Island, two skuas (unidentified species), were reported and confirmed positive for HPAI H5N1. The animals were collected at the Primavera Antarctic Base scientific station, at Cape Primavera, at 64°09’00 ″S 60°57′50″W. The result of this finding has not been published yet, but it was informed in the media. Additionally, a separate report informs suspected cases of HPAI in the Antarctic territory: Leon et al (2024) in their 2024 preprint, document suspected cases of HPAI affecting Adélie penguins and Antarctic shags at Red Rock Ridge, Antarctic region (67.18°W, 68.29°S). Due to their reliance on conventional PCR assays, these samples have been tentatively classified as suspected cases (León et al., 2024). Interestingly, these cases did not correspond to any observed mortality; all the animals appeared healthy, a finding that contradicts most of the existing results. All cases in the Antarctic region can be found at https://scar.org/library-data/avian-flu.

On the contrary, throughout the observation period, at the Fildes Peninsula, the Antarctic wildlife exhibited normal behaviors and no significant clinical signs indicative of HPAI. A significant number of samples were collected from marine mammals and seabirds. Moreover, the absence of unexpected mortality events and the negative results from real-time RT-PCR assays suggest that HPAI probably was not present in the wildlife at the Fildes Peninsula during the surveillance period, at least in the species with a large number of samples.

The lack of viral detection at Fildes Peninsula raises questions about the dynamics of HPAI in Antarctica. Previously, along the northern Chilean coast, the virus has rapidly propagated hundreds of kilometers weekly (Pardo-Roa et al., 2023; Ariyama et al., 2023). If we consider that the southern place with HPAI confirmed by October 8^th^, 2023, was Bird Island, South Georgia, it was reasonable to think the detection and further of HPAI H5N1 within two months in Antarctica. The lack of evident dissemination of the virus should be related to several factors, and the case observed at Bird Island can help. The first confirmed species at Bird Island were Brown skuas, and then successive events of mortality in marine mammals and seabirds were documented in the location, although the virus was confirmed only in Brown skuas and gulls (Bennison et al., 2023).

Furthermore, our observations indicate a decrease in the skua population at the site, which could align with the sightings of deceased individuals in other locations previously mentioned, as well as the confirmed cases at Primavera Cape and our case at James Ross Island. The noticeable decrease in skua populations at Fildes Peninsula could be related to HPAIV, where infected birds have died on their migratory routes and haveńt arrived in the breeding area. However, as there were many breeding seasons with comparable low numbers, other factors such as the impact of the late summer with of lot of snow at the normal time of breeding or a lag of marine food resources could play a more important role (Krietsch et al., 2016).

The confirmation of HPAI H5N1 in skua species in the other locations, as well as its apparent smaller population observed at the season at Fildes Peninsula, could suggest this species could present higher susceptibility of HPAI H5N1 compared to other Antarctic species. Other skua species such as Great skua have been confirmed positive for HPAI H5N1, and even a massive mortality event has been documented at Foula Island, United Kingdom (Furness et al., 2023; Lean et al., 2023). On the other hand, as top predators, and scavengers devoid of natural enemies, and with widely spaced nests due to territorial behavior, Antarctic skuas likely present a reduced risk for rapid virus dissemination between them. However, in some circumstances can be transmitted to other animals causing local outbreaks as those observed at Bird Island, South Georgia. Regarding elucidating the hypothesis, in the absence of massive mortalities in Antarctic territory, the study of skua species is mandatory not only for direct viral detection and sequencing but serology.

In addition, other seabird species can move between temperate, subantarctic, and Antarctic waters, often visiting the South American coast for foraging or overwintering, like Southern giant petrels (Petersen 2017, Finger 2023), or suspected to realize migratory movements during the non-breeding season, such as kelp gulls and snowy sheathbills.

These birds have been identified as potential vectors of infectious pathogens in this vulnerable ecosystem due to their scavenging habits, and previously documented roles as carriers of low-pathogenicity avian influenza viruses (LPAIV) (Barriga et al., 2016; de Souza Petersen et al., 2017; de Seixas et al., 2022). Kelp gulls, Snowy sheathbills, and Brown skuas are often mentioned as main vectors of the disease due to their short-range migratory capabilities (i.e.(Krasnobaev et al., 2018; Padilha et al., 2023). This complexity presents challenges in elucidating their potential contributions to disease dissemination.

However, no tracking study for Kelp gulls or Snowy Sheathbills has been published confirming migratory or dispersing movements, therefore the role of those species in the long-range dispersion of viruses is still to be confirmed. Also, Fur seals’ participation in HPAI dissemination cannot be discarded, since males migrate from South Georgia Island (March et al., 2021), a location with reported positive cases during the period of the study. The role of individuals with long-distance/transequatorial migration in the transmission of HPAI into Antarctica cannot be dismissed since they have been pivotal in the transcontinental dissemination of the HPAI virus from Europe to North America (Caliendo et al., 2022b) and from North America to South America (Leguia et al., 2023). Finally, further studies on population dynamics pre- and post-HPAI and serologic evaluation to measure antibody levels may be necessary to assess the population-wide consequences of HPAI H5N1.

The arrival of H5N1 in Antarctica poses a significant risk to wildlife. Outbreaks in South Africa, Chile, and Argentina have demonstrated high vulnerability of *Spheniscus* penguins (Molini et al., 2020; Pardo-Roa et al., 2023). However, to date, no mass mortality events associated with *Pygoscelis* penguins have been reported. On the other hand, a massive mortality event caused by HPAI H5N1 was recently reported in Argentina, in which hundreds of pups of Southern elephant seals died (Campagna et al., 2024).

During the surveillance period, we confirmed HPAI H5N1 in South polar skuas at James Ross Island, but no clinical signs attributable to HPAI H5N1 were observed at Fildes Peninsula. Based on our studies, the viral dynamic in the Antarctic seems to be slower compared to other regions, most likely with a low transmission rate. Further studies are necessary to evaluate the role of key species, such as skuas, in the dynamics of HPAI in Antarctica. Additionally, sequencing of positive cases is crucial for more detailed genetic characterization.

## Data Availability Statement

The data that support the findings of this study are available from the corresponding author, V.N., upon reasonable request.

## Acknowledgments

We thank the INACH personnel for supporting the sample collection and logistics. Thanks also go to the Czech Antarctic Infrastructure (CzechPolar2) at James Ross Island for the support provided during sample collection.

## Contributions

Conceptualization, M.G., G.M, and V.N.; methodology, B.B.L., B.B., L.K., and G.M; formal analysis, V.N.; investigation, N.E., G.M, M.G., and V.N.; resources, M.G., L.K., and V.N.; data curation, B.B.L.; writing—original draft preparation, B.B.L., B.B., L.K., G.M., C.B., and V.N.; writing—review and editing, C.B., and V.N.; project administration, V.N.; funding acquisition, M.G., L.K., and V.N. All authors have read and agreed to the published version of the manuscript.

## References

Ariyama, N., C. Pardo-Roa, G. Muñoz, C. Aguayo, C. Ávila, C. Mathieu, B. Brito, R. Medina, M. Johow, and V. Neira, 2023 (7. April): Emergence and rapid dissemination of highly pathogenic avian influenza virus H5N1 clade 2.3.4.4b in wild birds, Chile. 2023.04.07.535949, DOI: 10.1101/2023.04.07.535949. bioRxiv.

Barriga, G.P., D. Boric-Bargetto, M.C. San Martin, V. Neira, H. van Bakel, M. Thompsom, R. Tapia, D. Toro-Ascuy, L. Moreno, Y. Vasquez, M. Sallaberry, F. Torres-Pérez, D. González-Acuña, and R.A. Medina, 2016: Avian Influenza Virus H5 Strain with North American and Eurasian Lineage Genes in an Antarctic Penguin. Emerg Infect Dis 22, 2221–2223, DOI: 10.3201/eid2212.161076.

Bennison, A., A.M.P. Byrne, S.M. Reid, J.G. Lynton-Jenkins, B. Mollett, D.D. Sliva, J. Peers-Dent, K. Finlayson, R. Hall, F. Blockley, M. Blyth, M. Falchieri, Z. Fowler, E.M. Fitzcharles, I.H. Brown, J. James, and A.C. Banyard, 2023 (24. November): Detection and spread of high pathogenicity avian influenza virus H5N1 in the Antarctic Region. 2023.11.23.568045, DOI: 10.1101/2023.11.23.568045. bioRxiv.

Boulinier, T., 2023: Avian influenza spread and seabird movements between colonies. Trends in Ecology & Evolution 38, 391–395, DOI: 10.1016/j.tree.2023.02.002.

Braun, C., O. Mustafa, A. Nordt, S. Pfeiffer, and H.-U. Peter, 2012: Environmental monitoring and management proposals for the Fildes Region, King George Island, Antarctica. Polar Research DOI: 10.3402/polar.v31i0.18206.

British Antarctic Survey, 2024 (17. February): Additional cases of Avian Flu confirmed on South Georgia [Online] Available at https://www.bas.ac.uk/media-post/additional-cases-of-avian-flu-hpai-confirmed-on-south-georgia/ (accessed February 17, 2024).

Caliendo, V., N.S. Lewis, A. Pohlmann, S.R. Baillie, A.C. Banyard, M. Beer, I.H. Brown, R. a. M. Fouchier, R.D.E. Hansen, T.K. Lameris, A.S. Lang, S. Laurendeau, O. Lung, G. Robertson, H. van der Jeugd, T.N. Alkie, K. Thorup, M.L. van Toor, J. Waldenström, C. Yason, T. Kuiken, and Y. Berhane, 2022a: Transatlantic spread of highly pathogenic avian influenza H5N1 by wild birds from Europe to North America in 2021. Sci Rep 12, 11729, DOI: 10.1038/s41598-022-13447-z.

Caliendo, V., N.S. Lewis, A. Pohlmann, S.R. Baillie, A.C. Banyard, M. Beer, I.H. Brown, R. a. M. Fouchier, R.D.E. Hansen, T.K. Lameris, A.S. Lang, S. Laurendeau, O. Lung, G. Robertson, H. van der Jeugd, T.N. Alkie, K. Thorup, M.L. van Toor, J. Waldenström, C. Yason, T. Kuiken, and Y. Berhane, 2022b: Transatlantic spread of highly pathogenic avian influenza H5N1 by wild birds from Europe to North America in 2021. Sci Rep 12, 11729, DOI: 10.1038/s41598-022-13447-z.

Campagna, C., M. Uhart, V. Falabella, J. Campagna, V. Zavattieri, R.E.T. Vanstreels, and M.N. Lewis, 2024: Catastrophic mortality of southern elephant seals caused by H5N1 avian influenza. Marine Mammal Science 40, 322–325, DOI: 10.1111/mms.13101.

Charostad, J., M. Rezaei Zadeh Rukerd, S. Mahmoudvand, D. Bashash, S.M.A. Hashemi, M. Nakhaie, and K. Zandi, 2023: A comprehensive review of highly pathogenic avian influenza (HPAI) H5N1: An imminent threat at doorstep. Travel Medicine and Infectious Disease 55, 102638, DOI: 10.1016/j.tmaid.2023.102638.

de Seixas, M.M.M., J. de Araújo, S. Krauss, T. Fabrizio, D. Walker, T. Ometto, L. Matsumiya Thomazelli, R.E.T. Vanstreels, R.F. Hurtado, L. Krüger, R. Piuco, M.V. Petry, R.G. Webster, R.J. Webby, D.-H. Lee, D.H. Chung, H.L. Ferreira, and E.L. Durigon, 2022: H6N8 avian influenza virus in Antarctic seabirds demonstrates connectivity between South America and Antarctica. Transbound Emerg Dis 69, e3436–e3446, DOI: 10.1111/tbed.14728.

de Souza Petersen, E., J. de Araujo, L. Krüger, M.M. Seixas, T. Ometto, L.M. Thomazelli, D. Walker, E.L. Durigon, and M.V. Petry, 2017: First detection of avian influenza virus (H4N7) in Giant Petrel monitored by geolocators in the Antarctic region. Mar Biol 164, 62, DOI: 10.1007/s00227-017-3086-0.

Dewar, M., M. Wille, A. Gamble, R.E.T. Vanstreels, T. Bouliner, A. Smith, A. Varsani, N. Ratcliffe, J. Black, A. Lynnes, A. Barbosa, and T. Hart, 2023: The risk of highly pathogenic avian influenza in the Southern Ocean: a practical guide for operators and scientists interacting with wildlife. Antarctic Science 35, 407–414, DOI: 10.1017/S0954102023000342.

Egevang, C., I.J. Stenhouse, R.A. Phillips, A. Petersen, J.W. Fox, and J.R.D. Silk, 2010: Tracking of Arctic terns Sterna paradisaea reveals longest animal migration. Proceedings of the National Academy of Sciences 107, 2078–2081, DOI: 10.1073/pnas.0909493107.

Furness, R.W., S.C. Gear, K.C.J. Camphuysen, G. Tyler, D. de Silva, C.J. Warren, J. James, S.M. Reid, and A.C. Banyard, 2023: Environmental Samples Test Negative for Avian Influenza Virus H5N1 Four Months after Mass Mortality at A Seabird Colony. Pathogens 12, 584, DOI: 10.3390/pathogens12040584.

Gass Jr, J.D., R.J. Dusek, J.S. Hall, G.T. Hallgrimsson, H.P. Halldórsson, S.R. Vignisson, S.B. Ragnarsdottir, J.E. Jónsson, S. Krauss, S.-S. Wong, X.-F. Wan, S. Akter, S. Sreevatsan, N.S. Trovão, F.B. Nutter, J.A. Runstadler, and N.J. Hill, 2023: Global dissemination of influenza A virus is driven by wild bird migration through arctic and subarctic zones. Molecular Ecology 32, 198–213, DOI: 10.1111/mec.16738.

Hindell, M.A., C.R. McMahon, M.N. Bester, L. Boehme, D. Costa, M.A. Fedak, C. Guinet, L. Herraiz-Borreguero, R.G. Harcourt, L. Huckstadt, K.M. Kovacs, C. Lydersen, T. McIntyre, M. Muelbert, T. Patterson, F. Roquet, G. Williams, and J.-B. Charrassin, 2016: Circumpolar habitat use in the southern elephant seal: implications for foraging success and population trajectories. Ecosphere 7, e01213, DOI: 10.1002/ecs2.1213.

Kopp, M., H. Peter, O. Mustafa, S. Lisovski, M. Ritz, R. Phillips, and S. Hahn, 2011: South polar skuas from a single breeding population overwinter in different oceans though show similar migration patterns. Marine Ecology Progress Series 435, 263–267, DOI: 10.3354/meps09229.

Krasnobaev, A., G. ten Dam, S.P.J. van Leeuwen, L.S. Peck, and N.W. van den Brink, 2018: Persistent Organic Pollutants in two species of migratory birds from Rothera Point, Adelaide Island, Antarctica. Marine Pollution Bulletin 137, 113–118, DOI: 10.1016/j.marpolbul.2018.10.008.

Krietsch, J, S. Hahn, M. Kopp, R. Phillips, H. Peter, and S. Lisovski, 2017: Consistent variation in individual migration strategies of brown skuas. Mar. Ecol. Prog. Ser. 578, 213– 225, DOI: 10.3354/meps11932.

Krietsch, Johannes, J. Esefeld, C. Braun, S. Lisovski, and H.-U. Peter, 2016: Long-term dataset reveals declines in breeding success and high fluctuations in the number of breeding pairs in two skua species breeding on King George Island. Polar Biol 39, 573–582, DOI: 10.1007/s00300-015-1808-7.

Lane, J.V., J.W.E. Jeglinski, S. Avery-Gomm, E. Ballstaedt, A.C. Banyard, T. Barychka, I.H. Brown, B. Brugger, T.V. Burt, N. Careen, J.H.F. Castenschiold, S. Christensen-Dalsgaard, S. Clifford, S.M. Collins, E. Cunningham, J. Danielsen, F. Daunt, K.J.N. D’entremont, P. Doiron, S. Duffy, M.D. English, M. Falchieri, J. Giacinti, B. Gjerset, S. Granstad, D. Grémillet, M. Guillemette, G.T. Hallgrímsson, K.C. Hamer, S. Hammer, K. Harrison, J.D. Hart, C. Hatsell, R. Humpidge, J. James, A. Jenkinson, M. Jessopp, M.E.B. Jones, S. Lair, T. Lewis, A.A. Malinowska, A. McCluskie, G. McPhail, B. Moe, W.A. Montevecchi, G. Morgan, C. Nichol, C. Nisbet, B. Olsen, J. Provencher, P. Provost, A. Purdie, J.-F. Rail, G. Robertson, Y. Seyer, M. Sheddan, C. Soos, N. Stephens, H. Strøm, V. Svansson, T.D. Tierney, G. Tyler, T. Wade, S. Wanless, C.R.E. Ward, S.I. Wilhelm, S. Wischnewski, L.J. Wright, B. Zonfrillo, J. Matthiopoulos, and S.C. Votier n.d.: High pathogenicity avian influenza (H5N1) in Northern Gannets (Morus bassanus): Global spread, clinical signs and demographic consequences. Ibis **n/a**, DOI: 10.1111/ibi.13275.

Lean, F.Z.X., M. Falchieri, N. Furman, G. Tyler, C. Robinson, P. Holmes, S.M. Reid, A.C. Banyard, I.H. Brown, C. Man, and A. Núñez, 2023: Highly pathogenic avian influenza virus H5N1 infection in skua and gulls in the United Kingdom, 2022. Vet Pathol 3009858231217224, DOI: 10.1177/03009858231217224.

Leguia, M., A. Garcia-Glaessner, B. Muñoz-Saavedra, D. Juarez, P. Barrera, C. Calvo-Mac, J. Jara, W. Silva, K. Ploog, Lady Amaro, P. Colchao-Claux, M.M. Uhart, M.I. Nelson, and J. Lescano, 2023: Highly pathogenic avian influenza A (H5N1) in marine mammals and seabirds in Peru (preprint). Genomics.

León, F., C.L. Bohec, E.J. Pizarro, L. Baille, R. Cristofari, A. Houstin, D.P. Zitterbart, G. Barriga, E. Poulin, and J.A. Vianna, 2024 (18. March): Highly Pathogenic Avian Influenza A (H5N1) Suspected in penguins and shags on the Antarctic Peninsula and West Antarctic Coast. 2024.03.16.585360, DOI: 10.1101/2024.03.16.585360. bioRxiv.

March, D., M. Drago, M. Gazo, M. Parga, D. Rita, and L. Cardona, 2021: Winter distribution of juvenile and sub-adult male Antarctic fur seals (Arctocephalus gazella) along the western Antarctic Peninsula. Sci Rep 11, 22234, DOI: 10.1038/s41598-021-01700-w.

McCarthy, A.H., L.S. Peck, David C Aldridge, and de, 2022: Ship traffic connects Antarctica’s fragile coasts to worldwide ecosystems. Proceedings of the National Academy of Sciences 119, e2110303118, DOI: 10.1073/pnas.2110303118.

Molini, U., G. Aikukutu, J.-P. Roux, J. Kemper, C. Ntahonshikira, G. Marruchella, S. Khaiseb, G. Cattoli, and W.G. Dundon, 2020: Avian Influenza H5N8 Outbreak in African Penguins (Spheniscus demersus), Namibia, 2019. J Wildl Dis 56, 214–218.

Padilha, J.A., G.O. Carvalho, W. Espejo, A.R.L. Pessôa, L.S.T. Cunha, E.S. Costa, J.P.M. Torres, G. Lepoint, K. Das, and P.R. Dorneles, 2023: Trace elements in migratory species arriving to Antarctica according to their migration range. Marine Pollution Bulletin 188, 114693, DOI: 10.1016/j.marpolbul.2023.114693.

Pardo-Roa, C., M.I. Nelson, N. Ariyama, C. Aguayo, L.I. Almonacid, G. Munoz, C. Navarro, C. Avila, M. Ulloa, R. Reyes, E.F. Luppichini, C. Mathieu, R. Vergara, Á. González, C.G. González, H. Araya, J. Fernández, R. Fasce, M. Johow, R.A. Medina, and V. Neira, 2023: Cross-species transmission and PB2 mammalian adaptations of highly pathogenic avian influenza A/H5N1 viruses in Chile. bioRxiv 2023.06.30.547205, DOI: 10.1101/2023.06.30.547205.

Pertierra, L.R., K.A. Hughes, G.C. Vega, and M.Á. Olalla-Tárraga, 2017: High Resolution Spatial Mapping of Human Footprint across Antarctica and Its Implications for the Strategic Conservation of Avifauna. PLOS ONE 12, e0168280, DOI: 10.1371/journal.pone.0168280.

Prosser, D.J., J. Chen, C.A. Ahlstrom, A.B. Reeves, R.L. Poulson, J.D. Sullivan, D. McAuley, C.R. Callahan, P.C. McGowan, J. Bahl, D.E. Stallknecht, and A.M. Ramey, 2022: Maintenance and dissemination of avian-origin influenza A virus within the northern Atlantic Flyway of North America. PLOS Pathogens 18, e1010605, DOI: 10.1371/journal.ppat.1010605.

Sumner, M.D., S.J. Wotherspoon, and M.A. Hindell, 2009: Bayesian Estimation of Animal Movement from Archival and Satellite Tags. PLOS ONE 4, e7324, DOI: 10.1371/journal.pone.0007324.

Ulloa, M., A. Fernández, N. Ariyama, A. Colom-Rivero, C. Rivera, P. Nuñez, P. Sanhueza, M. Johow, H. Araya, J.C. Torres, P. Gomez, G. Muñoz, B. Agüero, R. Alegría, R. Medina, V. Neira, and E. Sierra, 2023: Mass mortality event in South American sea lions (Otaria flavescens) correlated to highly pathogenic avian influenza (HPAI) H5N1 outbreak in Chile. Vet Q 43, 1–10, DOI: 10.1080/01652176.2023.2265173.

WHO, 2011: WHO information for molecular diagnosis of influenza virus in humans - update. Who 1–38.

